# ESC - a comprehensive resource for SARS-CoV-2 immune escape variants

**DOI:** 10.1101/2021.02.18.431922

**Authors:** Mercy Rophina, Kavita Pandhare, Afra Shamnath, Mohamed Imran, Bani Jolly, Vinod Scaria

**Author notes:** **Address of the author for correspondence** Vinod Scaria, MBBS, PhD, Senior Scientist, G N Ramachandran Knowledge Centre for Genome Informatics, CSIR Institute of Genomics and Integrative Biology (CSIR-IGIB), South Campus, Mathura Road, New Delhi 110020. **Email of authors** Mercy Rophina; Kavita Pandhare; Afra Shamnath; Mohamed Imran; Bani Jolly.

## Abstract

Ever since the breakout of COVID-19 disease, ceaseless genomic research to inspect the epidemiology and evolution of the pathogen has been undertaken globally. Large scale viral genome sequencing and analysis have uncovered the functional impact of numerous genetic variants in disease pathogenesis and transmission. Emerging evidence of mutations in spike protein domains escaping antibody neutralization is reported. We have built a database with precise collation of manually curated variants in SARS-CoV-2 from literature with potential escape mechanisms from a range of neutralizing antibodies. This comprehensive repository encompasses a total of 5258 variants accounting for 2068 unique variants tested against 230 antibodies, patient convalescent plasma and vaccine breakthrough events. This resource enables the user to gain access to an extensive annotation of SARS-CoV-2 escape variants which would contribute to exploring and understanding the underlying mechanisms of immune response against the pathogen. The resource is available at http://clingen.igib.res.in/esc/

**GRAPHICAL ABSTRACT:** 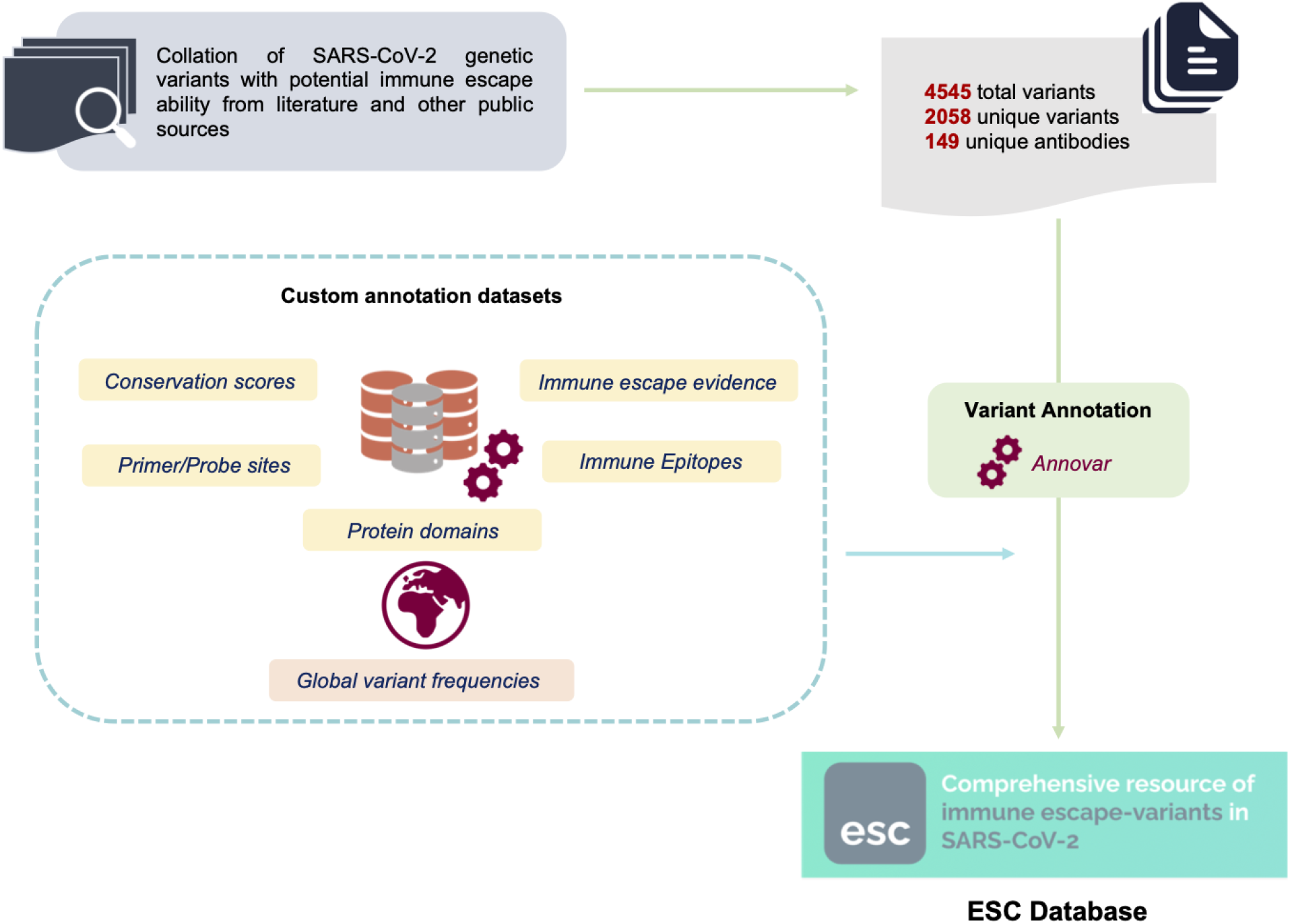

## INTRODUCTION

Genomic approaches have been instrumental in understanding the origin and evolution of SARS-CoV-2, the causative agent for the COVID-19 pandemic (1). Availability of the genome sequence of one of the earliest SARS-CoV-2 genomes from Wuhan province (2) and high throughput approaches to resequence and analyse viral genomes have facilitated the availability of numerous open genomic data sharing initiatives by the researchers worldwide. Pioneering public sources like GenBank (3) and Global Initiative on Sharing all Influenza Data (GISAID) (4) provide access to systematically organized genomes of SARS-CoV-2. The China National GeneBank DataBase (CNGBdb) (5), Genome Warehouse (GWH) (6) and Virus Pathogen Resource (ViPR) (7) are few other resources which provide access to viral genomes and perform analyses on phylogeny, sequence similarity and genomic variants.

There has been a significant interest in recent times in understanding the functional impact of genetic variants in SARS-CoV-2 apart from exploring the genetic epidemiology. The variant D614G present in spike protein has been one the earliest and prominent examples with potential implications associated with the infectivity of the virus (8). Studies explaining the possible impact of SARS-CoV-2 variants in diagnostic primers and probes have augmented the importance of analysing the variations and their underlying role in disease pathogenesis (9). Various resources have been made available to help comprehend the virus better and also to understand its evolution. Public sources exclusively documenting functionally relevant SARS-CoV-2 variants based on literature evidence are also available (10).

With the advent of therapies including monoclonal antibodies, convalescent plasma as well as the recent availability of vaccines, interest in genetic variants which could affect the efficacy of such modalities of therapy has accelerated. The targeting of spike proteins by broad-neutralizing antibodies against SARS-CoV-2 offers a potential means of treating and preventing further infections of COVID-19 (11). Evidence on immunodominant epitopes with significantly higher response rates have also been reported (12). Antibody response to SARS-CoV-2 is one of the key immune responses which is actively being pursued to develop therapeutic strategies as well as vaccines (13). The recent months have seen enormous research into the structural and molecular architecture of the interactions between the spike protein in SARS-CoV-2 and antibodies. Studies have also provided insights into the genetic variants which could confer partial or complete resistance to antibodies (14) as well as panels of convalescent plasma. With vaccines being widely available, the evidence on the effect of genetic variants on efficacy of vaccines is also emerging (15)

The lack of a systematic effort to compile genetic variants in SARS-CoV-2 associated with immune escape motivated us to compile the information in a relevant, searchable and accessible format. Towards this goal, we systematically evaluated publications for evidence on immune escape associated with genetic variants in SARS-CoV-2 and created a database named as ESC. User-friendly web interface is made available to retrieve information on immune escape variants as well as their extensive functional annotations. To the best of our knowledge, this is the first most comprehensive resource for immune escape variants for SARS-CoV-2. The resource can be accessed online at http://clingen.igib.res.in/esc/.

## MATERIALS AND METHODS

### Data and Search Strategy

Genetic variants in the SARS-CoV-2 genome and evidence suggesting association with immune escape were systematically catalogued. A significant number of variants were associated with escape or resistance to a range of neutralizing and monoclonal antibodies, while a subset was associated with resistance to convalescent plasma. The data was compiled by manual curation of literature available from peer-reviewed publications and preprints. Literature reports with relevant information on antibody escape variants were retrieved from sources including PubMed, LitCovid, Google Scholar and preprint servers. The reports were systematically checked for details pertaining to the variation, antibodies tested and experimental methods followed in the study. In addition, the variants were systematically categorized based on experimental validation and computational prediction. Collated data was organized in a pre-formatted template based on their protein positions. This comprehensive compendium was used for further functional annotations.

### Variant Information and Annotations

The variant information and annotations were retrieved from annotation tables for individual features using ANNOVAR (16). Variant annotations broadly included genic features like the variant type and functional annotations related to deleteriousness and evolutionary conservation. Information on protein domains and immune epitopes was compiled and customized from various public sources. Variant sites reported to be potentially problematic including homoplasic regions, sites with recurrent sequencing errors and hypermutable sites were also labelled to enable quality check of the mutation site. Variants mapping back to sites of potential SARS-CoV-2 diagnostic primers/probes were also annotated.

### Compilation of B-cell and T-cell epitope data

Details on B-cell and T-cell epitopes spanning the protein residues of SARS-CoV-2 were retrieved from Immune Epitope Database and Analysis Resource (IEDB) (17). All epitopes of SARS-CoV-2 (IEDB ID : 2697049) against human hosts with reported positive or negative assays and any type of MHC restriction were used for analysis. Epitope information pertaining to each amino acid residue including the epitope type (linear/discontinuous), epitope sequence with corresponding start and end positions and IEDB identifiers were systematically mapped back and documented.

### Antibody details and annotation

Information pertaining to the list of antibodies associated with escape mechanisms was retrieved from available public sources. Compiled antibodies were systematically mapped back to the AntiBodies Chemically Defined (ABCD) database which provides integrated information regarding the antibodies along with its corresponding antigens and protein cross links to fetch unique antibody identifiers (18, 19).

### Database and Web Interface

The back-end of the web interface was implemented using Apache web server and MongoDB v3.4.10 in order to provide a user-friendly interface for variant search. The JavaScript Object Notation (JSON) file format was used to systematically store the data. PHP 7.0, AngularJS, HTML, Bootstrap 4, and CSS were used to code the web interface for querying. Highcharts javascript library was also used for improved presentation and interactivity. A Beacon API has been created using the PHP programming language. The ESC Beacon API v1.0.0 is a read-only API with specifications written in OpenAPI. It uses JSON in requests and responses and standard HTTPS for information transfer. The Beacon API has one endpoint: /beacon?Variant; query interface is provided by the beacon endpoint.

## RESULTS & DISCUSSION

### Repository of SARS-CoV-2 escape variants

We compiled a total of 5258 variant entries from over 60 articles which studied SARS-CoV-2 variants and their effect on immune escape. This included a total of 2068 unique variants mapping to spike protein, ORF1ab and ORF3a. Out of the total unique variants, 2060 variants mapped to the gene coding for spike protein with potential immune escape mechanisms elucidated through experimental evidence as well as computational predictions. The remaining 8 variants were found in ORF1ab and ORF3a genes, out of which, 3 were reported to confer potential epitope loss. The compiled list of variants was found associated with 230 unique SARS-CoV-2 antibodies and patient polyclonal sera. A handful of SARS-CoV-2 variations associated with vaccine breakthrough events have also been documented. A brief comparison of the curations in the ESC database with other publicly available resources is summarized in **Supplementary Figure 1** and **Supplementary Table 1a and 1b**. Functional consequences of the variants were mapped from a total of 22 unique custom generated annotation datasets precisely including deleteriousness and conservation score predictions, protein domains and immune epitopes using ANNOVAR. The data features used in the study are summarised in **Supplementary Table 2**.

### Antibody association mapping

By scanning through the spike protein residues and their associations with SARS-CoV-2 neutralizing and monoclonal antibodies, we were able to compile the exact count of antibodies reported to have potential associations with the residues. From our analysis we observed that spike protein residues ranging from 350 to 500 amino acid positions exhibited potential antibody associations with the possibility of immune escape against at least one antibody. A total of 22 hotspot residues (140,144,246,248,346,417,439,444,445,446,450,452,453,455,475,477,484,485,486,490,49 3,501) were found to possess immune evasion capability against >10 monoclonal antibodies. A schematic representation of the number of antibodies associated with spike protein residues along with their domain annotations is shown in **Figure 1**. The cumulative frequencies of spike mutation sites associated with immune escape against >5 mAbs in the receptor binding domain is shown in **Figure 2**. Systematic categorization of mutation residues along with their localization in spike protein is depicted in **Figure 3**.

**Figure 1.**
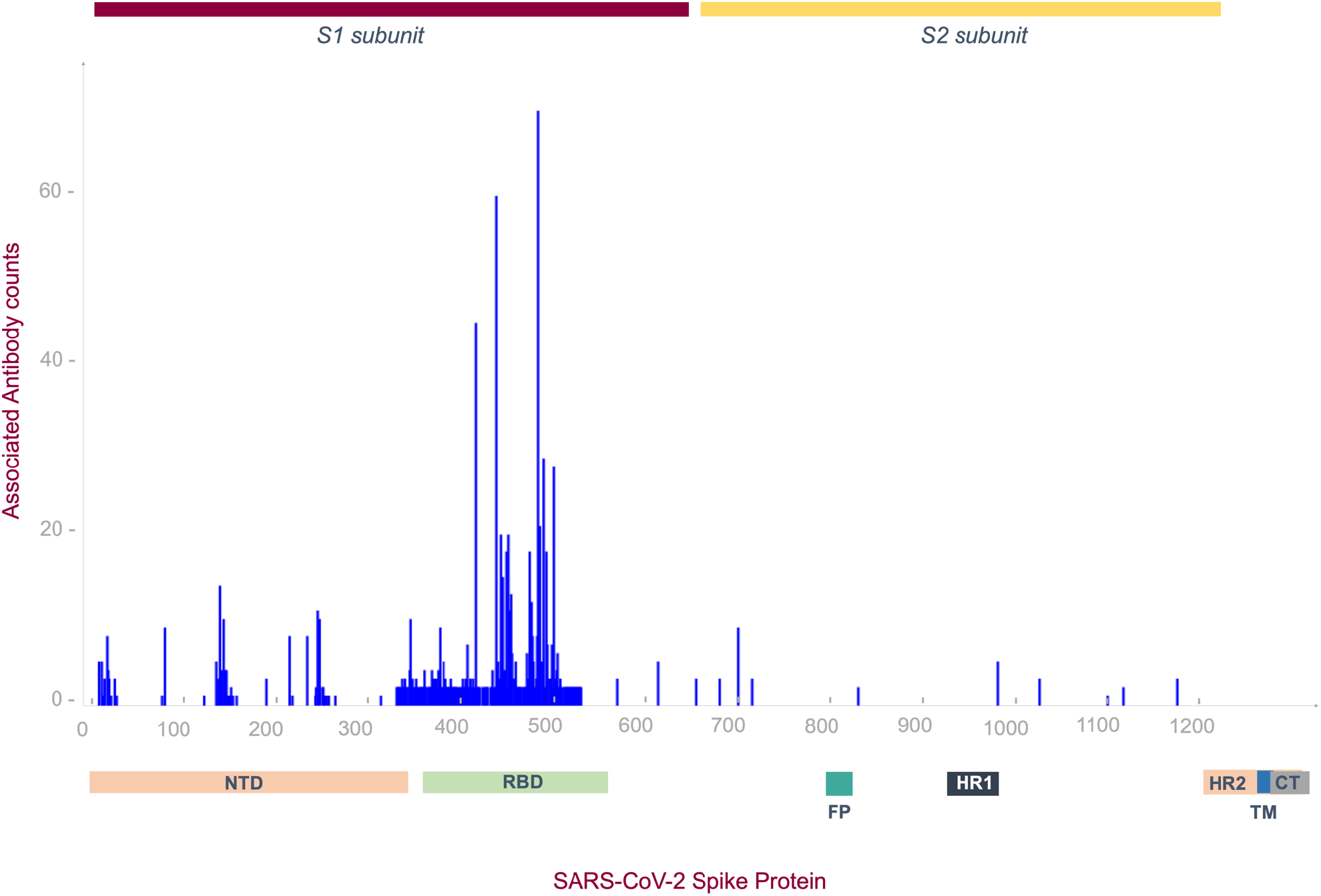
Illustration of mutation - antibody associations along the spike protein residues. Number of unique antibodies associated with potential immune evasion at each spike protein residue is marked along with domain annotations.

**Figure 2.**
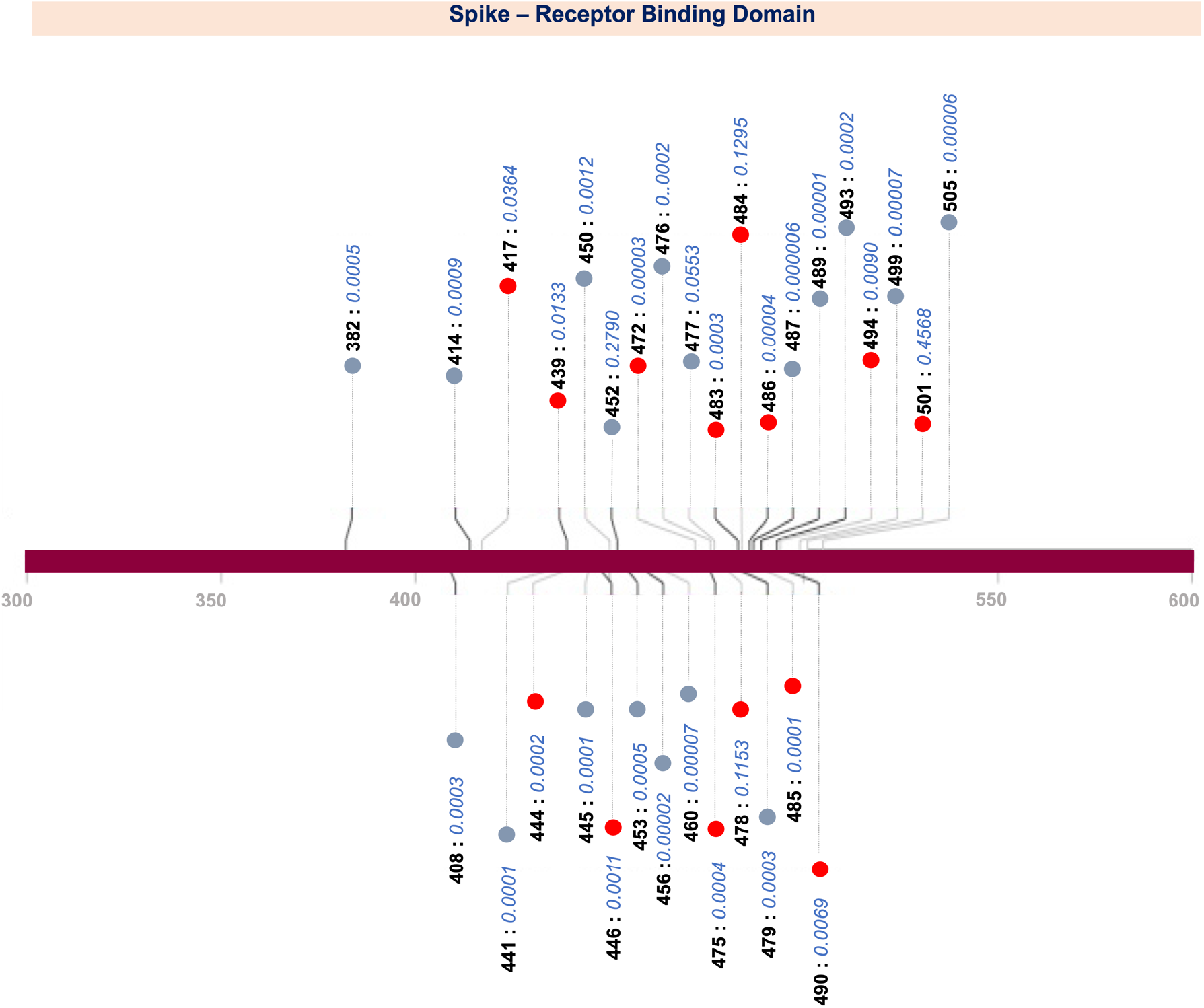
Distribution of mutation sites associated with >5 mAbs in the Receptor binding domain of SARS-CoV-2 spike protein. Variant sites with potential impact on neutralization of human polyclonal sera are represented in red. Cumulative frequencies of variants at RBD sites are mentioned alongside.

**Figure 3.**
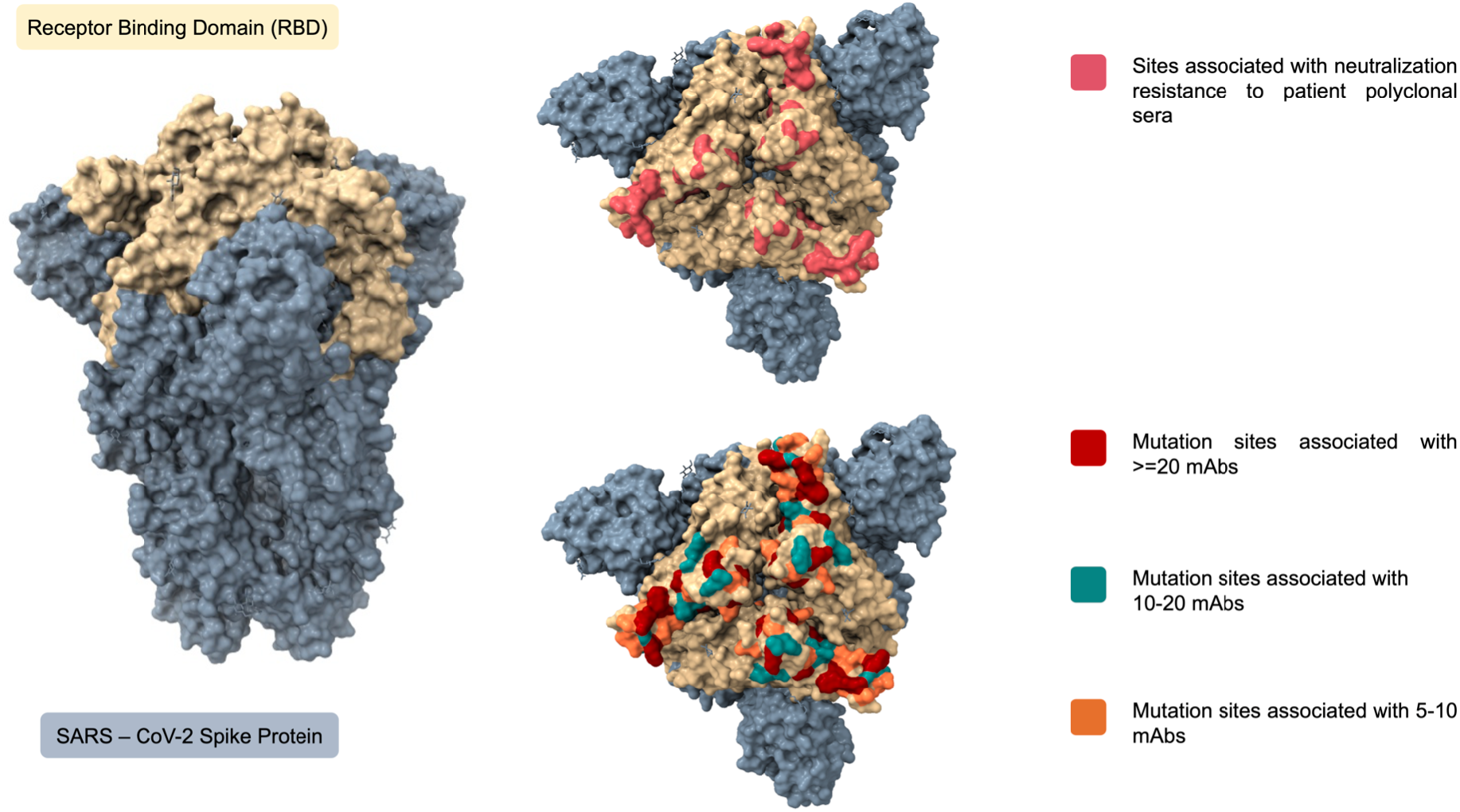
Receptor Binding Domain of spike protein with mutation hotspot sites associated with decreased neutralization against patient polyclonal sera and monoclonal antibodies.

### Overview of B cell epitopes and immunodominant epitope regions

With the aim of mapping back the known B and T cell epitopes encompassing the variant compendium, SARS-CoV-2 epitope details were extracted. There were a total of 310 and 472 experimentally validated B cell and T cell epitopes respectively. This precisely included 263 linear and 47 discontinuous B cell epitopes in spike protein. Reported B cell and T cell epitope information was mapped back to residues possessing antibody escape mutations which provided a brief insight on the potential impact of these variations in immune recognition and responses.

### Database features

The database offers a user-friendly interface enabling the users to search for variants based on their amino acid change, gene name or the antibody name as per the specified format. The search query returns a list of matching results, whose complete functional annotations can be viewed by clicking on the displayed elements. The resource provides a list of annotation features for each variant precisely organized into 8 major sections namely Variant details, Antibody details, Variants of Concern/Interest, Protein domain details, Epitope details, Functional annotation, Literature Evidence and Variant frequency. **Figure 4a and b** portrays the query search and the result display section of the resource

**Figure 4.**
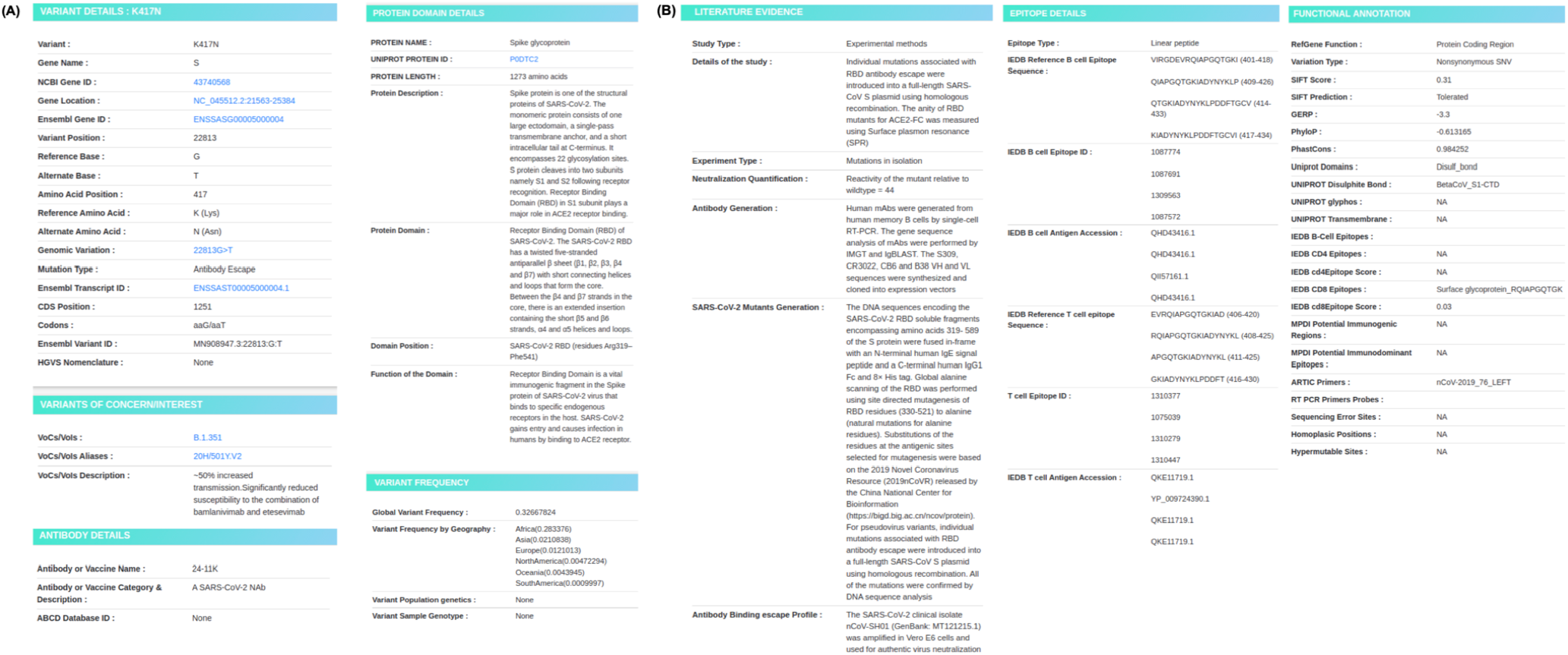
Panel illustrating the query search and display features in ESC database.

Basic details pertaining to the variant like the amino acid change, genomic coordinates and the variant type have been enlisted in the Variant Details section. Information on the associated neutralizing antibodies and their identifiers are provided in the Antibody details section. Domain and epitope details section exclusively comprises details on the protein domain, epitopes reported to span the protein residue through experimental validations. Computationally predicted functional annotations on deleteriousness from SIFT (20), evolutionary conservation scores provided by PhastCons (21), GERP (22) and PhyloP (23) are included in the Functional annotation section. This section also enlists protein domain information retrieved from UniProt and immune epitopes documented from IEDB (17, 24), UCSC and predictions from different software packages (B cells-BepiPred 2.0, CD4-IEDB Tepitool, CD8-NetMHCpan4). Annotations of potential error prone sites including sites of sequencing errors, homoplasic and hypermutable regions (https://virological.org/t/issues-with-sars-cov-2-sequencing-data/473) and diagnostic primer/probe sites are also mapped. Extensive evidence from literature curation including the methods of the study, neutralization quantification profiles and details of antibody/mutant generation are summarized in the literature evidence section. Variant frequency section exclusively summarizes the estimated frequencies of the variant on a global scale as well as by its geography. In addition, characteristic mutations of VoCs and VoIs have also been annotated with brief descriptions in the Variants of Concern/Interest section.

## CONCLUSIONS

With evidence emerging on genetic variants in SARS-CoV-2 associated with resistance to monoclonal antibodies and convalescent plasma using *in-vitro* assays, unique insights into the structural and functional mechanisms whereby the pathogen could evolve and evade antibodies have become possible. These insights could have enormous implications in efficacy of vaccines currently being used as well as under trials. One of the recent studies has reported the impact of a few immune escape variants on the efficacy of vaccines (25). It is expected that similar studies would be extended for a wider number of variants as well as vaccines. In order to keep pace with rapid discoveries regarding SARS-CoV-2 escape variations and mechanisms, the database and the associated Github repository is being updated every month from peer reviewed publications and pre-print articles with complete annotations. We therefore foresee that the ESC resource would be a central resource to enable such studies and provide a ready reference to the emerging evidence on immune escape.

## Supporting information

ESC Supplementary Materials

## DATA AVAILABILITY

The completed data curated from various literature sources are collated and made available for access and bulk download at https://github.com/mercywilliams160896/ESC_COVID19 An API for ESC is also made available for ease of access to data. Example search : https://clingen.igib.res.in/esc/api/beacon?Variant=A475V. A detailed overview of The ESC Beacon API v1.0.0 has been documented and linked to the webpage duly for user interests.

## AUTHOR’S CONTRIBUTIONS

MR, AS and MI systematically curated data for the database. KP designed the database. BJ provided annotation details and helped in fact checks. VS conceived and designed the project. All authors approved the final manuscript.

## ACKNOWLEDGEMENTS

Authors acknowledge Anjali Bajaj for her constructive comments and suggestions.

## FUNDING

This work was funded by The Council of Scientific and Industrial Research (CSIR), India through CODEST grant. The funders have no role in the analysis or decision to publish.

## CONFLICT OF INTEREST

None declared.

## Notes

https://clingen.igib.res.in/esc/

### Competing Interest Statement

The authors have declared no competing interest.

### Summary of Updates

The updated features of ESC database has been highlighted in the revised version of the manuscript

