## Supplementary material for "ESC - a comprehensive resource for SARS-CoV-2 immune escape variants": ESC Supplementary Materials

### SUPPLEMENTARY DATASETS

**Supplementary Figure 1.** A brief comparison of the SARS-CoV-2 immune escape variant details from publicly available resources

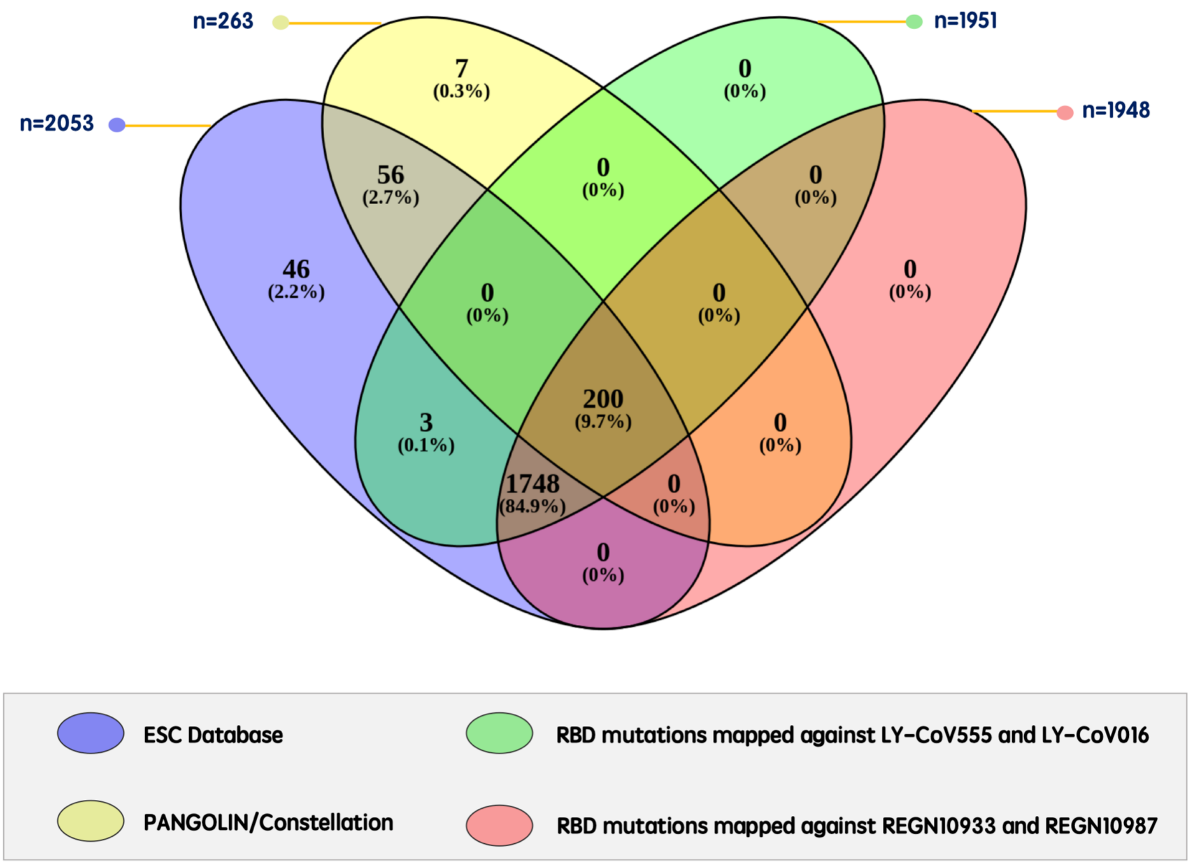

**Supplementary Table 1a.** Summary of publicly available resources with systematic collation of SARS-CoV-2 immune evasion variants

| S.No | Resource/Repository | Contents and highlights | Total entries count | Resource/Repository Link |
|---|---|---|---|---|
| 1. | PANGOLIN/Constellation | ***List of immune escape variants*** in Spike protein along with the ***associated antibodies*** | **263** | https://github.com/cov-lineages/constellations/blob/main/constellations/data/escape_list.csv (not available now) |
| 2. | Complete map of SARS-CoV-2 RBD mutations that escape the monoclonal antibody LY-CoV555 and its cocktail with LY-CoV016 | List of experimentally validated ***immune escape fractions*** for mutations sites in ***Spike RBD*** against ***LY-CoV555, LY-CoV016 and cocktail of LY-CoV555 and LY-CoV016*** | **1951** | https://github.com/jbloomlab/SARS-CoV-2-RBD_MAP_LY-CoV555/blob/main/results/supp_data/LY_cocktail_raw_data.csv |
| 3. | Prospective mapping of viral mutations that escape antibodies used to treat COVID-19 | List of experimentally validated ***immune escape fractions*** for mutations sites in ***Spike RBD*** against ***REGN10933, REGN10987 and cocktail of REGN10933 and REGN10987*** | **1948** | https://github.com/jbloomlab/SARS-CoV-2-RBD_MAP_clinical_Abs/blob/main/results/supp_data/REGN_and_LY-CoV016_raw_data.csv |
| 4. | Coronavirus Antiviral and Resistance Database | List of variations and their corresponding escape fractions comprehensively collected from literature sources | **All reported VoCs and VoIs with isolated mutations also** | https://covdb.stanford.edu/page/susceptibility-data/ |

**Supplementary Table 1b.** A complete summary of the highlights and contents of the resources and repositories of SARS-CoV-2 immune escape variants

| **Resource/Repository** | **Access** | **Interface options** | **Variant annotations** |
|---|---|---|---|
| PANGOLIN Constellation | Publicly available | No search interface | List of SARS-CoV-2 escape mutations with associated mAbs |
| RBD mutations mapped against LY-CoV555 and LY-CoV016 | Publicly available | No search interface | Estimates of total and maximum escape fractions for RBD escape mutation sites against Ab cocktails |
| RBD mutations mapped against REGN10933 and REGN10987 | Publicly available | No search interface | Estimates of total and maximum escape fractions for RBD escape mutation sites against Ab cocktails |
| Coronavirus Antiviral and Resistance Database | Publicly available | User friendly search options available | Provides fold reductions in neutralization against a range of monoclonal antibodies and patient sera |
| ESC database | Publicly available | User friendly search interface available | Annotations on<br>• VOCs/VoIs<br>• Protein Domains<br>• B cell and T cell Epitopes<br>• Associated mAbs,patient sera or vaccines<br>• Conservation sites<br>• Prediction of deleteriousness<br>• Sites spanning diagnostic primers/probes and potential error prone regions<br>• Global and geographical variant frequencies (including a graphical representation)<br><br>Extensive literature curation on<br>• Type of study<br>• Experimental details<br>• Details of neutralization escape efficacy |

| | | | |
|---|---|---|---|
| | | | • Generation of monoclonal antibodies and mutant viruses<br>Other available linkouts<br>• User friendly linkouts to GISAID and CoVDB have been provided for extensive search of the variants |

**Supplementary Table 2.** List of custom annotation features used for ANNOVAR annotations

| Custom Datasets |
|---|
| **Variation Type** |
| *Non synonymous variant* |
| *Synonymous variant* |
| **SIFT_Score** |
| **SIFT_Prediction (***Deleterious/Tolerable)* |
| **GERP** |
| **PhyloP** |
| **PhastCons** |
| **Uniprot_Domains** |
| **UNIPROT_Disulphite_Bond** |
| **UNIPROT_glyphos** |
| **UNIPROT_Transmembrane** |
| **IEDB_B-Cell_Epitopes** |
| **IEDB_CD4_Epitopes** |
| **IEDB_cd4Epitope_Score** |
| **IEDB_CD8_Epitopes** |
| **IEDB_cd8Epitope_Score** |
| **MPDI_Potential_Immunogenic_Regions** |
| **MPDI_Potential_Immunodominant_Epitopes** |
| **ARTIC_Primers** |
| **RT-PCR_Primers/Probes** |
| **Sequencing_Error_Sites** |
| **Homoplasic_Positions** |
| **Hypermutable_Sites** |
